# How Diverse are the Mountain karst Forests of Mexico?

**DOI:** 10.1101/2023.03.01.530643

**Authors:** María Eugenia Molina-Paniagua, Pablo Hendrigo Alves de Melo, Santiago Ramírez-Barahona, Andrés Ernesto Ortiz-Rodriguez, Alexandre K. Monro, Carlos Manuel Burelo-Ramos, Héctor Gómez-Domínguez

**Affiliations:** Posgrado, División Académica de Ciencias Biológicas, Universidad Juárez Autónoma de Tabasco, Villahermosa, Tabasco, México, 86150; Departamento de Botánica, Instituto de Biología, UNAM, Ciudad Universitaria, Apartado Postal 70-367, 04510, Ciudad de México, México; UNESP-Universidade Estadual Paulista “Julio de Mesquita Filho”, Río Claro, São Paulo, SP, Brazil, 13506-900; Americas Team, The Herbarium, Royal Botanic Gardens Kew, UK, TW9 3AB; Herbario UJAT, División Académica de Ciencias Biológicas, Universidad Juárez Autónoma de Tabasco, Villahermosa, Tabasco, México, 86150; Senda sustentable, A.C., Tuxtla Gutiérrez, Chiapas, México

## Abstract

Tropical forests on karstic relief (karst forest) are among the most species-rich biomes. These forests play pivotal roles as global climate regulators and for human wellbeing. Their long-term conservation could be central to global climate mitigation and biodiversity conservation. In Mexico, karst landscapes occupy 20% of the total surface and are distributed mainly in the southeast of the country, along the eastern slope, and in the Yucatan Peninsula. Within each of these areas, the following types of karst occur: coastal karst, plain karst, hill karst, and low-, medium and high-mountain karst. Mountain karst forests cover 2.07% of Mexico’s surface and are covered by tropical rainforests, montane cloud forests, and tropical deciduous forests. It is probably one of the most diverse biomes in Mexico. However, the Mountain karst forest of Mexico has received little attention, and very little is known about it. Here, we evaluate the vascular plant species richness within the mountain karst forests of Mexico. We assembled the first, largest and most comprehensive datasets of Mexican, mountain karst forest species, from different public databases (CONABIO, GBIF, IBdata-UNAM), which included a critical review of all data. The families, genera and species present within the mountain karst forest of Mexico were compiled. Taxa that best characterize the forest of Mexico were identified based on their spatial correlation to this biome. Also, the conservation status of each of them was determined. We explored biodiversity patterns, identifying areas with the highest species richness, endemism centers, and areas of relatively low sampling intensity. We found that within the mountain karst forest of Mexico there are representatives of 11,771 vascular plant species (253 families and 2,254 genera), ca. 50 % of the Mexican flora. We identified 372 species endemic to these forests. According to preliminary IUCN red list criteria, 2,477 species are under some category of conservation risk, of which 456 (3.8 %) are endangered. Most of the Mexican karst forests have been extensively explored and six allopatric, species-rich areas were identified. Compared to other regions in the world, the mountain karst forest of Mexico is one of the most diverse biomes. It contains more species than some entire montane systems in Mexico such as Sierra Madre Oriental, and Sierra Madre del Sur. Also, the mountain karst forest of Mexico is most diverse than similar forests of South America and Asia, even if considering the effect of different sampling areas. The fact that mountain karst forest is covered by some of the most species-rich Mexican forests, in addition to being embedded in areas of high biotic diversity in Mexico, probably contributes to its great floristic diversity. Thus, the mountain karst forest of Mexico is an important source of diversity and shelters a large percentage of the Mexican flora.

## Introduction

Limestone karst forests are one of the most diverse biomes worldwide^1^. This high diversity is attributed to their soil conditions, high heterogeneity of microhabitats, and their archipelago-like distribution^2–7^. Plant lineages that have colonised and diversified in these forests are specialized and characterised by small and generally disjunct distribution ranges, and with notorious morphological innovations^6,8–11^.

In Mexico, karst landscapes occupy 20% of the total surface (~391 700 km^2^) and is distributed mainly in the southeast of the country (Chiapas, Guerrero and Oaxaca), along the eastern slope, and in the Yucatan Peninsula^12–14^. Within each of these areas the following types of karst occur: coastal karst, plain karst, hill karst, and low-, medium and high-mountain karst^15^. Each type is determined by its structure, elevation, and the climatic conditions that drove the karstification process^13,14^. Mountain karst (named hereafter as Mountain karst forest) are distributed discontinuously through the Sierra Madre Oriental, from the southwest of Tamaulipas to southern Veracruz, in the northeastern portion of Oaxaca, in southern Tabasco and several regions of the state of Chiapas (Fig 1). In contrast, coastal karst, plain karst, and hill karst are best represented and almost restricted to the Yucatan Peninsula (=Karst of the Yucatan Peninsula^13,14^ or platform plains and hills karst ^15^).

**Figure 1.**
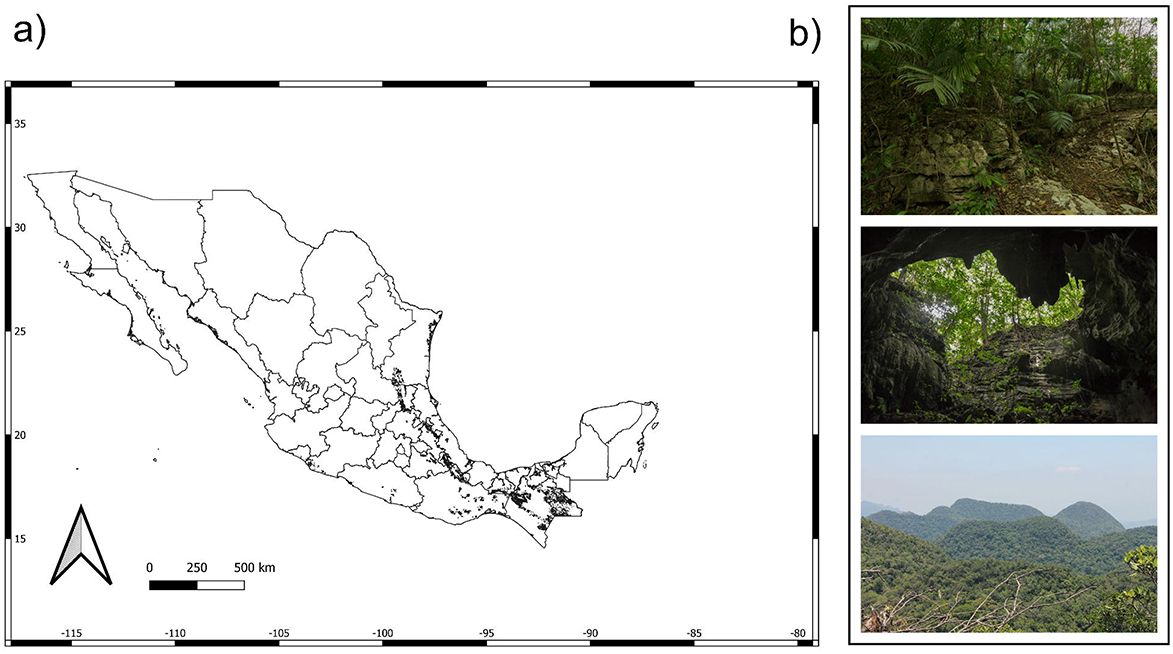
Mountain karst forest of Mexico. A) it is limited to small, archipelago like-distributed remnants along the mountain systems of western (Guerrero and Michoacán), eastern (Puebla, Querétaro, San Luis Potosí, Tamaulipas, and Veracruz), and southern Mexico (Chiapas, Oaxaca, and Tabasco), and occupying an area of 40,759.87 km^2^ (2.07% of Mexico surface). B) Most of the Mountain-limestone karst forest area is occupied by the tropical rain forest. In these forests, the chasms, cliffs, caves and limestone soils characterize the landscape.

Mountain karst forests cover 2.07% of Mexico’s surface, and harbor part of some species-rich forests in the country, such as tropical rain forest, montane cloud forest, and deciduous forest^13,14^; Fig 1). Despite that, studies focused on mountain karst forests are scarce (eg Wendt^2^; Vázquez-Torres^16^; Gonzales-Medrano and Hernández-Mejía^17^; Pérez-Farrera et al. ^10^; Ramírez-Marcial et al.^18^; Kovarik and Beynen^19^), and estimates of its total diversity are based on very low sampling effort and so likely to be underestimated. Moreover, mountain karst forests encompass biodiversity hotspots where new species are described steadily, yet species extinction is increasing rapidly^22^ with the loss of habitats as the main factor. The availability of large and public databases, containing historical and contemporary taxonomic information, collected over several decades in many parts of the world and often accompanied by collection, ecological and geographic information^20,21^, offers an opportunity to estimate the plant diversity of this biome.

Here, we assembled the first comprehensive datasets of the Mexican, mountain karst forest species, from different public databases, accompanied by the critical review of all data. We sought to determine the number of species restricted to the mountain karst forest of Mexico and the conservation status of each of them.

We also explored biodiversity patterns, identifying species richness areas, endemism centers, and little-explored regions. Specifically, by answering the following questions: 1) How diverse is the mountain karst forest of Mexico? 2) What are the spatial patterns of species richness distribution? 3) Where are the centers of endemism? 4) How many species are restricted to mountain karst forest? and, 5) What is the conservation status of each of them?

## Methods

### Data acquisition

Fig 2 summarizes the workflow used to obtain and validate the names that underpin the vascular plant dataset for Mexico, and which was used to extract occurrence data for mountain karst forest of Mexico:

**Figure. 2.**
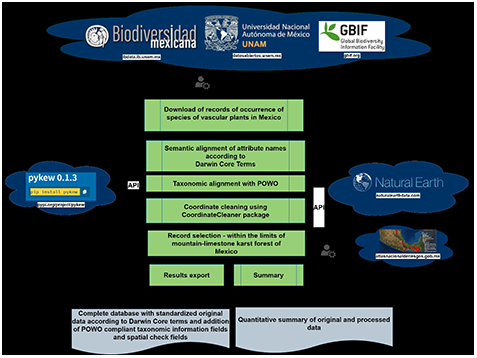
Workflow for composing vascular plants dataset for Mexico and extracting flora from mountain karst forest of Mexico.

We obtained 7,655,611 records of the Mexican vascular plant flora. These were sourced from, 1) the Comisión Nacional para el Conocimiento y Uso de la Biodiversidad^23^; 47.7% of total records), 2) the Global Biodiversity Information Facility (GBIF^24^; 36.8 % of total records; (https://doi.org/10.15468/dl.af62vr), and 3) the web system of the scientific collections of the Institute of Biology-UNAM^25^ <u>[IBdata v3 «Helia Bravo Hollis» (https://www.ibdata.abaco3.org)</u>; 15.5 % of total records]. To facilitate content comparison and manipulation between data sources, a semantic alignment of attribute names was undertaken according to Darwin Core Terms. This was followed by a harmonization of taxonomic names using Kew’s Plants of the World database^26^ using the *pykew* 0.1.3 ^27^ R-package. We discarded records that were not identified to the rank of species. Synonyms, duplicate specimens, and invalid, unmatched or missing names were also discarded. This resulted in 6, 492, 388 records, an attrition of 16% for Mexican vascular plants. We also discarded records lacking accurate geolocations according to the following criteria: a) non-numeric and not available coordinates (9.2% of total records); b) zero longitude or latitude (10 records flagged); c) coordinates outside the reference landmass (23.5% of records): d) coordinates in vicinity of Country and province centroids (0.5% of total records); e) coordinates in vicinity of country capitals (0.3 % of total records); f) Records inside urban areas (3.8 % of total records); g) records in the vicinity of biodiversity institutions (0.1% of total records), and h) duplicated records (16.3% of total records).. We did so using *CoordinateCleaner R*-package^28^. This resulted in a further attrition of 14% and a dataset of 5,820,489 occurrence records representing 21,659 species of Mexican vascular plants^29^. We extracted the mountain karst forest species from the above dataset using the shape file in Fig 1, where duplicate collections were eliminated based on the combination of cell (0.08 degrees), year, month, day and scientific name of collections, resulted in a first karst vascular plant dataset comprising of 184,804 occurrence records.

### Floristic characterization of mountain-limestone karst forest of Mexico and species affinity to karst

The families, genera and species present within the mountain karst forest of Mexico were compiled. Taxa that best characterize the mountain karst forest of Mexico were identified based on their spatial correlation to this biome. Specifically, we count the number of ‘karstic’ cells (0.5-degree latitude/longitude) that each species occupies (area of occupancy), by homogenizing the sampling effort to one record per cell, and in cases of combined cells (karstic, non-karstic), 1 record considered for each category. Thus, three types of spatial correlation were identified, 1) Mexico karst endemic species, whose total number of grid cells are exclusive to mountain karst forest of Mexico; 2) karst associated species, defined as those where more than half of a species’ grid cells occur within Mexican mountain karst forest; and, 3) non-karst species, defined as species where more than half of their grid cells occurs outside of Mexican mountain forest.

Species-richness and levels of endemism in Mexico was compared to karst forest biomes for which comparable information is available. This included karst forest in Brazil ^1^, Guizhou, China^30^, and the Malayan Peninsula^31^. To compare the species richness in areas of different sizes, a taxonomic biodiversity index was used. This calculates the number of observed species divided by the natural logarithm of the area in km2 (IB = E/lnA), where E is the number of recorded species and A the area^32^. Species-richness and endemism of the mountain karst forest of Mexico, was also compared to non-karst areas surroundings the study area, such as the Sierra Madre Oriental (SMO^33,34^) and the Sierra Madre del Sur (SMS^35, 36^).

### Biodiversity distribution patterns and the conservation status of species

Mountain-limestone karst forest of Mexico was divided into 89 grid cells (0.5-degree latitude/longitude grid size) for mapping species richness, endemism and conservation status. Each cell covered an area of approximately 256 km^2^, a medium-sized scale that avoids the effect of sampling artifacts, such as the occurrence of artificially empty grid squares ^37,38^. To assess whether the Mountain karst forest of Mexico is an area of high endemism, the weighted endemism (WE)^37^ and corrected endemism (CWE)^38^ were estimated using Biodiverse v.1.1 software^39^. We define endemic species as those restricted to the Mountain karst forest (all georeferenced records distributed exclusively within this biome). For each species within the Mountain karst forest of Mexico, we undertake a preliminary assessment of its conservation status by calculating the extent of occurrence (EOO) and the area of occupancy (AOO) using occurrence data, the function *iucn. eval* implemented in the *conR*, R-package^40^, and applying the IUCN Red List Categories and criteria. Within each grid cell, we separately measured and mapped the number of species, the endemic species, and the number of endangered species. We plotted this information and the resulting patterns (as maps) were analysed by overlaying layers for each parameter, enabling us to detect those areas with the highest species-richness, but also those regions that host most endemic and threatened species based on preliminary assessments. Finally, the maps were generated and projected in QGIS version 3.22.1-Białowieża^41^ and Infomap Bioregions^42^.

## Results

### Floristic characterization and species mountain karst forest affinity

A total of 11,771 species of vascular plants were recorded within the mountain karst forest of Mexico, distributed in 253 families and 2,254 genera. The 14 most diverse families in terms of number of species (more than 200 species) are presented in Table 1, which contribute more than 50% of all recorded species. The Piperaceae, Polypodiaceae and Orchidaceae families stands out among all because 91%, 78% and 62% of their Mexican species, respectively, were recorded within the mountain karst forest. Table 2 shows the plant families that characterize the forests studied, among them Gesneriaceae, Melastomataceae, Dioscoreaceae, Araceae, Violaceae, Urticaceae, Arecaceae, and Piperaceae, are the families with the highest proportion of their records distributed exclusively within the mountain karst forest of Mexico. The 10 genera with the highest number of species within the mountain-limestone karst forest are *Piper* (119 species), *Salvia* (111), *Peperomia* (103), *Ipomoea* (101), *Solanum* (100), *Tillandsia* (100), *Euphorbia* (95), *Epidendrum* (81), *Senna* (86), and Mimosa (78).

**Table 1.** The most diverse vascular plant families [more than 200 species within the mountain karst forest of Mexico (MLKF)]. The percentage of species with respect to the total number of species present in Mexico (^26,29^) is shown in parentheses.

| Family | Species in Mexico | Species within the MLKF |
| --- | --- | --- |
| Piperaceae | 245 | 222 (91%) |
| Polypodiaceae | 293 | 230 (78%) |
| Orchidaceae | 1213 | 760 (62%) |
| Solanaceae | 407 | 240 (59%) |
| Poaceae | 1047 | 606 (57%) |
| Acanthaceae | 385 | 220 (57%) |
| Fabaceae | 1903 | 1087 (57%) |
| Rubiaceae | 707 | 396 (55%) |
| Malvaceae | 527 | 284 (53%) |
| Apocynaceae | 418 | 213 (50%) |
| Cyperaceae | 416 | 210 (50%) |
| Euphorbiaceae | 714 | 340 (47%) |
| Lamiaceae | 601 | 229 (38%) |
| Asteraceae | 3057 | 1081 (35%) |

**Table 2.** Most common plant families with the highest proportion of records distributed exclusively within the mountain karst forest (MLKF) of Mexico.

| Families | Total records | Exclusive MLKF records (%) |
| --- | --- | --- |
| Gesneriaceae | 1283 | 14.06 |
| Melastomataceae | 2432 | 12.64 |
| Dioscoreaceae | 1069 | 12.53 |
| Araceae | 4944 | 12.24 |
| Violaceae | 735 | 12.24 |
| Urticaceae | 1440 | 11.97 |
| Arecaceae | 2648 | 11.68 |
| Piperaceae | 3200 | 11.22 |
| Begoniaceae | 1437 | 11.03 |
| Orchidaceae | 13364 | 10.80 |
| Primulaceae | 2264 | 10.68 |
| Marantaceae | 689 | 10.17 |
| Lauraceae | 4115 | 10.10 |
| Myrtaceae | 2521 | 9.73 |
| Araliaceae | 1816 | 9.72 |
| Meliaceae | 2576 | 9.05 |
| Bromeliaceae | 7751 | 8.92 |
| Smilacaceae | 1056 | 8.44 |
| Annonaceae | 1238 | 8.07 |
| Sapotaceae | 1469 | 7.96 |
| Phyllanthaceae | 902 | 7.69 |
| Rubiaceae | 9749 | 7.54 |
| Celastraceae | 1561 | 7.36 |
| Moraceae | 4392 | 7.20 |
| Loranthaceae | 728 | 7.03 |
| Cannaceae | 921 | 6.95 |
| Malpighiaceae | 3919 | 6.88 |
| Passifloraceae | 3352 | 6.57 |
| Phytolaccaceae | 1444 | 6.57 |
| Acanthaceae | 7499 | 6.38 |
| Gentianaceae | 1120 | 6.28 |
| Caricaceae | 1149 | 6.09 |
| Iridaceae | 2433 | 6.07 |
| Salicaceae | 2928 | 6.04 |
| Polygalaceae | 1135 | 6.03 |
| Campanulaceae | 2290 | 5.96 |
| Burseraceae | 8152 | 5.93 |
| Vitaceae | 1453 | 5.86 |
| Crassulaceae | 3407 | 5.86 |
| Petiveriaceae | 1297 | 5.80 |
| Lythraceae | 3038 | 5.70 |
| Malvaceae | 19919 | 5.66 |
| Bixaceae | 694 | 5.48 |
| Sapindaceae | 4147 | 5.36 |
| Apocynaceae | 17736 | 5.35 |
| Oleaceae | 818 | 5.32 |
| Combretaceae | 2465 | 5.29 |
| Bignoniaceae | 6716 | 5.21 |
| Rosaceae | 2876 | 5.10 |

We identified 372 endemic species (3.16% of the total species recorded in the mountain-limestone karst forest of Mexico), which are distributed in 84 families and 235 genera. The families with the largest number of endemic species are Orchidaceae (56 species), Fabaceae (26 species), Asteraceae (25 species), Piperaceae (19 species), Poaceae (13 species), Araceae (12 species), Aspleniaceae (12 species), Acanthaceae (10 species) and Polypodiaceae (10 especies). For other 731 species between 50% and 90% of their records are distributed exclusively within the mountain-limestone karst forest of Mexico (the mostly karstic species), whilst for the majority of the species recorded within this biome, 10,668 species, more than 50% of their records are present out the mountain-limestone karst forest of Mexico (the non-karstic species) (Fig 3). The proportion of records out the mountain karst forest, increases as a function of a higher number of total records in Mexico for each species (r^2^= 0.99, pval= < 2.2e-16). While 84% of the non-karstic species are very well represented in the collections and known from more than 20 localities, species endemic to the mountain-limestone karst forest are those known from a few scientific collections (mean= 2, max=17, min= 1).

**Figure 3.**
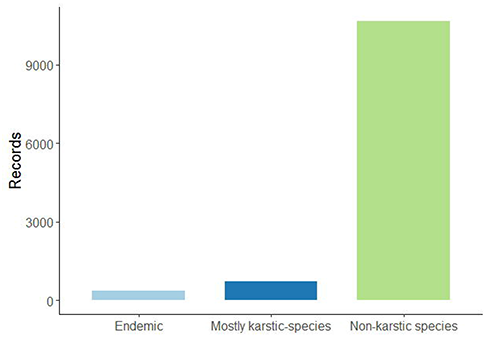
Proportion of records by species found out the mountain karst forest boundaries.

The taxonomic diversity index shows that the limestone karst forest of Mexico, is one of the most diverse compared to similar biomes in South America and Asia (Table 3).

**Table 3.** Taxonomic diversity index calculated for the mountain karst forest of Mexico and form similar biomes around the world. Limestone karst forest of Brazil, limestone karst forest of Guizhou, China, Malayan limestone karst forest Southeast Asia.

| Karst areas | Area (km<sup>2</sup>) | Species | Endemic species | Species/area | Taxonomic Biodiversity index |
| --- | --- | --- | --- | --- | --- |
| Brazil | 318,126 | 9,592 | 468 | 0.0302 | 757 |
| China | 170,000 | 7,505 | 171 | 0.0441 | 623 |
| Southeast Asia | 260 | 1,216 | 130 | 4.6769 | 219 |
| **Mexico** | **40,759** | **11,771** | **372** | **0.2586** | **993** |

### Biodiversity distribution pattern and conservation status of species

Species richness (the total number of recorded species in a cell) increased with the sampling effort (Pearson’s correlation= r^2^= 0.97, pval= < 2.2e-16). Although 38% of the grid cells within the limestone karst forest of Mexico have been poorly sampled [less than the median number of records (622) expected per cell), most of the forest regions (more than 60% of the grid cells analysed) recorded between 622 and 24,975 records. Moreover, major sampling efforts are not focused on a particular region and are rather randomly distributed within the study area (Fig S1). Accordingly, six allopatric cell-areas can be considered as species-rich: 1) the forests of Central Chiapas (Berriozabal-Chiapa de Corzo, 3,899 species), 2) the Zongolica region in Veracruz (Orizaba-Huatusco 2,574), extending up the Uxpanapan region in Veracruz (Uxpanapa, 1,755 species), 3) the forests of the highlands of Chiapas (Comitán-La Trinitaria, 2,124 species), 4) the Lacandon forest of Chiapas (Ocosingo, 1,985 species), 5) the Chimalapas region in Oaxaca (Chimalapas, 1,535), and 6) the Huasteca region between Queretaro, Hidalgo and San Luis Potosí (Cuichapa, 1531 species) (Fig 4).

**Figure 4.**
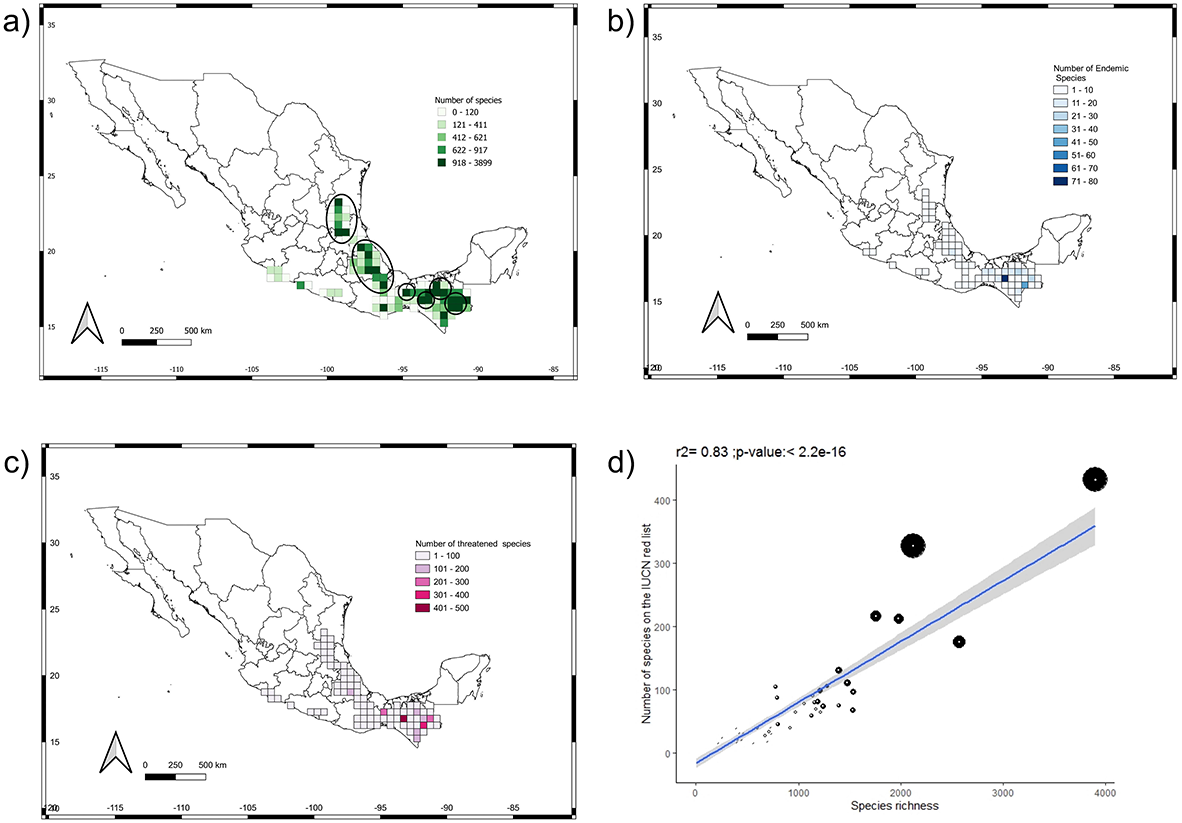
Species richness, endemism, and endangered species of the mountain karst forest of Mexico. A) Distribution of species richness. B) Distribution of endemism. C) Distribution of endangered species. D) Pearson’s correlation analysis between the species richness and the number of endangered species per cell, the size of the circles represents the number of endemic species. In each map, the species-rich areas, with more endemism and with more endangered species are circled (1) Central Chiapas, 2) the Zongolica and the Uxpanapa region, 3) Highlands of Chiapas, 4) the Lacandon forest, 5) Chimalapas region, 6) the Huasteca region.

We also identified 10 areas of endemism with 9 or more endemic species (Fig 4), distributed mainly in the south of Mexico, Chiapas, Oaxaca and Veracruz, but also with some regions identified in the east of Mexico, in Veracruz on the border with Puebla (the Zongolica region).

According to the IUCN red list^40^, there are 2,477 species in some category of risk (21% of the limestone karst forest species; Table 4). While in the Norma official Mexicana^43^, 376 endangered species (3.19% of the limestone karst forest species) are listed (Table 4). On the other hand, the tentative conservation status assessment (using the *ConR* package), suggests that at least 2,704 species (22.9% of the karst species) could be under some category of risk (Table 4, Fig 4).

**Table 4.** Species in, or potencially in a risk category according to the NOM-059 (Mexico) or IUCN category according to criterio B.

| Status | Nom_059 | In Red List | Putative in red list |
| --- | --- | --- | --- |
| LC o NT |  | 2,021 |  |
| VU | 136 | 186 | 1166 |
| EN | 147 | 221 | 1164 |
| CR | 91 | 45 | 374 |
| EW | 1 | 4 | □ |

Lastly, our overlay of richness, endemism, and endangered species layers revealed that the most species-rich regions contain a high number of endangered species, many of them endemic, being these priority areas for conservation (Fig 4). The above is critical considering that only a small portion, 2% of the limestone karst forest, is protected in some Mexican natural reserve (Fig S2).

## Discussion

In this study, we determined the number of species inhabiting the mountain-limestone karst forest of Mexico and the conservation status of each of them. We explored biodiversity patterns, identifying species richness areas, endemism centers, and little-explored regions:

### How diverse is the mountain-limestone karst forest of Mexico?

In Mexico, a wealth of 23,314 species of native vascular plants, 297 families and 2,854 genera have been estimated^29^. The species richness present in the limestone karst forest of Mexico (11,771 species, 2,254 genera, and 253 families) constitutes 50.49% of the Mexican plant species, 78.98% of Mexican genera, and 85.19% of the plant families. The fact that mountain karst forest is covered by some of the most species-rich Mexican forests, in addition to being embedded in areas of high biotic diversity in Mexico, probably contributes to its great floristic diversity. Contrary to expectations, the most diverse plant families in Mexico (eg Asteraceae, Rubiaceae, Lamiaceae^29^) are not the most species-rich families within the limestone karst forest of Mexico (Table 1), nor characterize the forests studied (Table 2). Instead, Gesneriaceae, Melastomataceae, Dioscoreaceae, Araceae, Violaceae, Urticaceae, Arecaceae, and Piperaceae, characterize the limestone karst forest of Mexico since are those plant families with the highest number of records distributed exclusively within this biome (Table 2). Among them, the Piperaceae family stands out with more than 90% of its species present in the limestone karst forest of Mexico (Table 1). Interestingly, this study revealed that there are plant lineages such as Elatinaceae, Nelumbonaceae, Frankeniaceae, Juncaginaceae, Nitrariaceae, Setchellanthaceae, Simmondsiaceae that are not favored by limestone karst soils and rather their diversity is concentrated in other biomes (coastal, aquatic, floodable, marshy or xeric habits).

We identified 372 species endemics to the mountain karst forest of Mexico where Orchidaceae, Fabaceae, Asteraceae, Piperaceae, Poaceae, Araceae, Aspleniaceae, Acanthaceae and are the plant families with more endemic species. The first three families are very diverse and widely distributed in Mexico^29^, their diversity within the limestone karst forest is not the highest, but the species that inhabit this biome tend to be endemic. A study in the Sierra Madre Oriental of Mexico supports the aforementioned and many genera of Asteraceae with a high number of endemic species are restricted to calcareous soils^34^. Similary, in Orchidaceae, a study in Asia^44^ suggests that the presence of many limestone karst forest-endemic species of this family, is a generalized pattern. For Araceae and Piperaceae, our results indicate that the diversity of both families is favored by their presence in karstic soils. Furthermore, from the total endemic species of Mexico (11,610 species/ ~50% of the Mexican flora^29^, 1, 424 (11%) have representatives within the limestone karst forest of Mexico, of which 3.2% (372 species) are restricted to this biome. Accordingly, the mountain karst forest of Mexico is an important source of diversity and shelters a large percentage of the Mexican flora.

Compared to other regions of Mexico and the world, the limestone karst forest (40, 759 km^2^, 11,771 species) is one of the most diverse biomes. It contains more species than some entire montane regions in Mexico such as Sierra Madre Oriental (220,151 km^2^, 6,981 species^34^), and Sierra Madre del Sur (143,447 km^2^, 7,016 species^35^). The limestone karst forest of Mexico is most diverse than similar forests of South America and Asia, even if considering the effect of different sampling areas (Table 3). Interestingly, our study revealed that the Neotropical karst forest (Brazil and Mexico) are more taxonomically diverse than the forests of Asia. However, Asian forests have a higher proportion of plant endemism than Neotropical forests (Table 3). Thus, it is probable that in the Neotropics the forest assemblages are phylogenetically more heterogeneous compared to Asia, but the speciation processes (species radiation) directly associated with limestone karst habitats are more frequent in Asia than in America^45–48^. This must be taken with caution as it is probably a generalized response that may not apply to particular families and genera in Mexico or neotropics as whole.

On the other hand, families with more endemic species within the mountain karst forest of Mexico are those with mainly herbaceous habits, such as Piperaceae, Araceae, Orchidaceae and Asteraceae. This is probably associated with the difficulty that rocky soils represent for the establishment of trees and shrubs. It is probable, then, that shallow soils with little organic matter and water favour the growth of herbaceous plants and limit the development of arboreal species^5^. Studies in herbs show that changes in genome size are associated with their presence in these habitats^46,47^.

### Biodiversity distribution pattern

Some authors based on the floristic pattern of particular regions^49,2^ considered the mountain karst forest of Mexico as centers of diversity and endemism, as a result of floristic refuges during periods of past climatic fluctuations. Our results support that hypothesis. Limestone karst forest of Mexico is characterized by a high number of species, many of them restricted to this biome, and with the richness and high endemism regions allopatrically distributed throughout the study region (Fig 4). A pattern associated with the presence of floristic refuges^2^. Six regions are here identified as areas of high species diversity and endemism, of which the Uxpanapan and the Zongolica regions, and central Chiapas have been proposed as floristic refuge areas (Fig S2)^2,49^. Our study suggests that in Mexico the limestone karst forest concentrate a high number of endemism, considering the georeferenced records of the species out and in the limestone karst forest (Fig 4). Assuming that in the rest of Mexico there is a greater sampling effort, the endemic and high diversity areas within the karst forest represent important areas for the preservation of Mexican flora.

In general, most high species richness areas are confined to topographically, ecologically and evolutionarily more complex regions of Mexico, which also form part of the biotic transition zone between the Holarctic and Neotropical regions of America. Thus, the mountain karst forest of Mexico develops on mountain systems, corridors and valleys, which, together with a complex climatic history, have driven the evolutionary history of its biota.

### Conservation status

Our study shows that the limestone karst forest of Mexico harbor many species with a restricted distribution range and that are therefore at extinction risk. The most diverse and high areas of endemism are also those with the most endangered species (Fig 4). These regions are undoubtedly a priority for the conservation of the Mexican limestone karst forest and for the Mexican flora as a whole. However, only 2% of the limestone karst forest are protected by natural reserves (Fig S3). Considering the georeferenced records in and out the limestone karst forest, more than 8, 000 species (more than 80% of the species recorded) could be under some risk category according to the criteria established by the IUCN (Table 3). If least concern species are not considered, more than twice of the species currently assessed by IUCN may need protection. This without considering that a much smaller number of species is currently protected by Mexican law (Table 3).

### Experimental error sources

Species richness and sampling effort are correlated (Fig S4). However, more than 60% of the forest studied shows high sampling values and the areas of high richness and endemism are randomly distributed. In addition, most of the cells analyzed fell below the regression line including two of the most species-rich sites, suggesting that indeed the forest has been relatively well sampled. The fact that we observed a high degree of correlation between species richness and sampling effort suggests that species-richness in species-poor sites was underestimated by collectors, possibly due to its geographical proximity to areas traditionally well explored or recognized as species-rich sites^1^.

Concerning the usefulness of public databases, our datasets gathers geographic information for 21659 species of vascular plants, a number very similar to the number of species estimated for the flora of Mexico [24,360^50^; 23,314^29^]. We obtained a total of 7, 655, 611 records from three public data sources, in order of contribution CONABIO, GBIF and IB data. After filtering, as detailed in the methods section, 76% of the records originally obtained from the three datasets were used in the final analyses. Poorly georeferenced data, duplicate geo-references, and duplicate collections, were the most frequently deleted records. Thus, the distribution pattern of species richness in Mexico based on our database is consistent with the pattern described in previous studies^29,50^ (Fig S5).

With respect to the taxonomic uncertainty, we standardize scientific names based on POWO^26^. From the scientific names originally obtained (56,302 species), 51% were retained as they were considered valid according to POWO. As detailed in the methods section, this cleaning also included the filtering of determinations above the species level, synonyms, duplicate collections, and missing names. Even so, our dataset probably includes “misdeterminations”, which directly influence the number of species estimated for Mexico and for the Mexican mountain karst forest. “Misdeterminations” have a complex origin, some probable factors are the little experience of botanists on the taxonomy of the species, the absence of local specialists, the absence of taxonomic keys or local and regional monographs, and of course, the species concept used (taxonomic issues). Taxonomic issues is ultimately due to the evolutionary nature of species, something hard to solve completely by standardization^51^. Interestingly, our species richness estimate and their distribution patterns for Mexico are quite similar to those inferred in other independent studies.

## Acknowledgments

We would like to thank Nelly Jimenez Pérez, Eduardo Ruiz Sánchez, German Carnevalli, Yuyini Licona Vera and anonymous reviewers, for providing many useful comments on the manuscript. We also wish to thank Christopher Davidson and Sharon Davidson for their support through the project “KARSTBIO: Biodiversity, Evolution and Conservation of the tropical rain forest on karstic zones of Latin America”. The first author is supported by a doctoral scholarship from CONACyT (212531). This work constitutes partial fulfillment of MEMP’s doctorate in Ecology and tropical ecosystem management at Universidad Juárez Autónoma de Tabasco (UJAT).

## Author contributions

MEMP, AEOR conceived the ideas, MEMP, SRB, AEOR and PHAM analyzed the data, MEMP and AEOR wrote the main manuscript text, AM, CMBR and HGD reviewed the manuscript.

## Supporting information

**S1 Fig**. Sampling effort distribution within the mountain karst forest of Mexico

**S2 Fig**. Species richness distribution within the mountain karst forest, the colored polygons indicate the location of the floristic refuges in Mexico according to Wendt (1989) and Toledo (1976).

**S3 Fig**. Mountain karst forest distribution and location of the main protected natural areas in Mexico

**S4 Fig**. Pearson’s correlation plot between species richness and sampling effort per cell (50 × 50).

**S5 Fig**. Species richness distribution of vascular plant in Mexico.

