## Supplementary figures and images for "How Diverse are the Mountain karst Forests of Mexico?"

### Fig S1

#### Slide 1
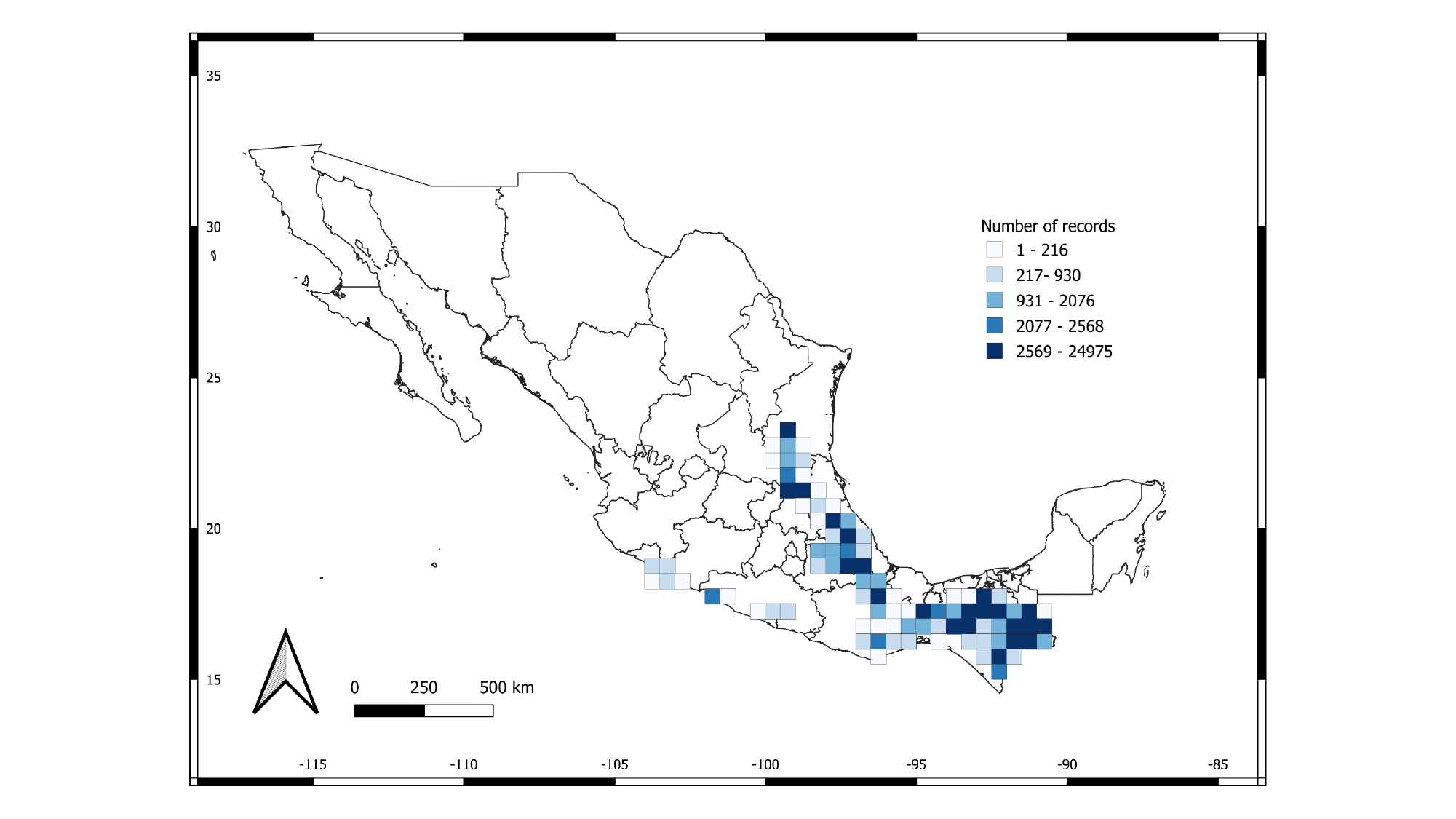

### Fig S2

#### Slide 1
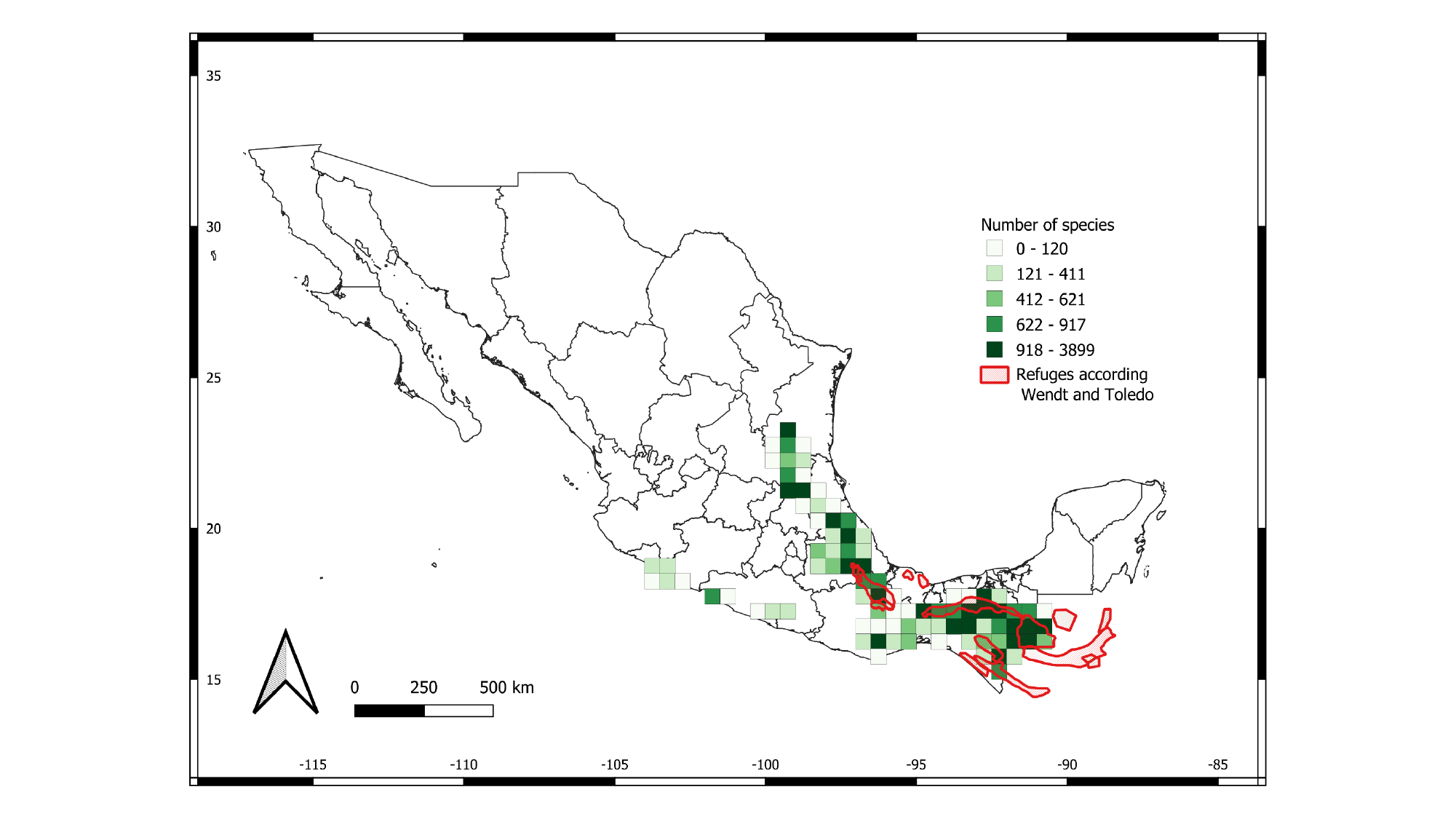

### Fig S3

#### Slide 1
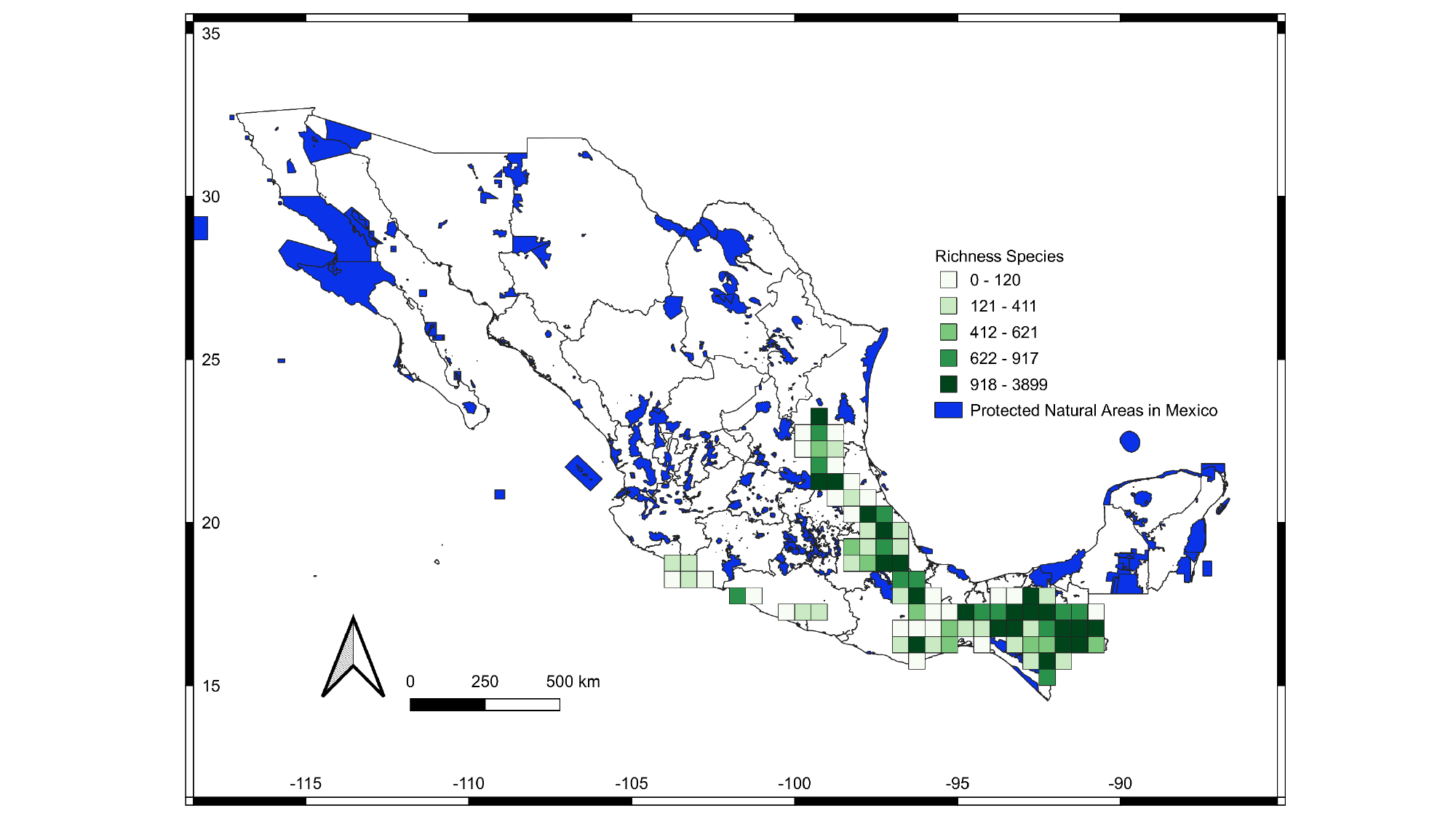

### Fig S4

#### Slide 1
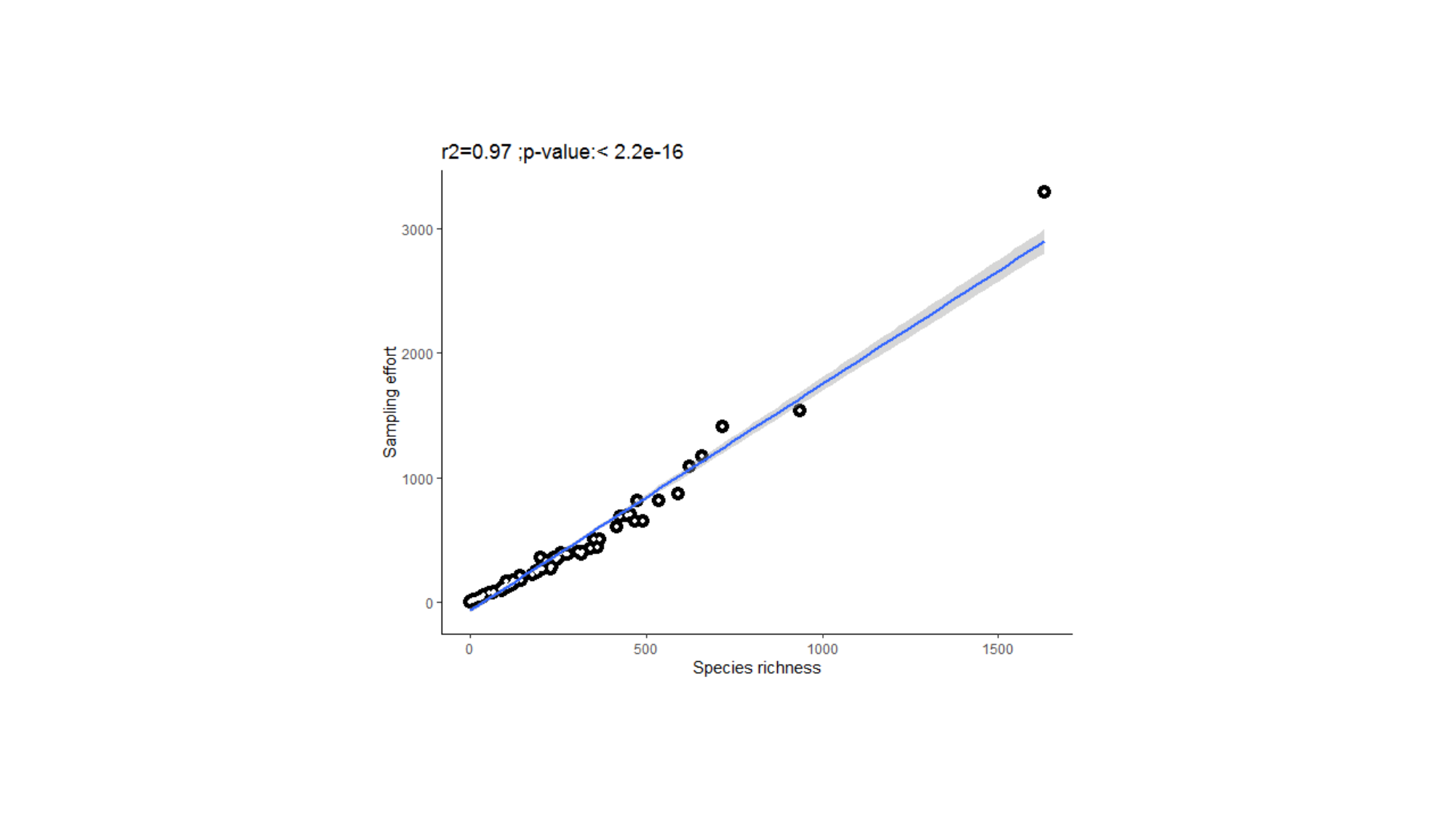

### Fig S5

#### Slide 1
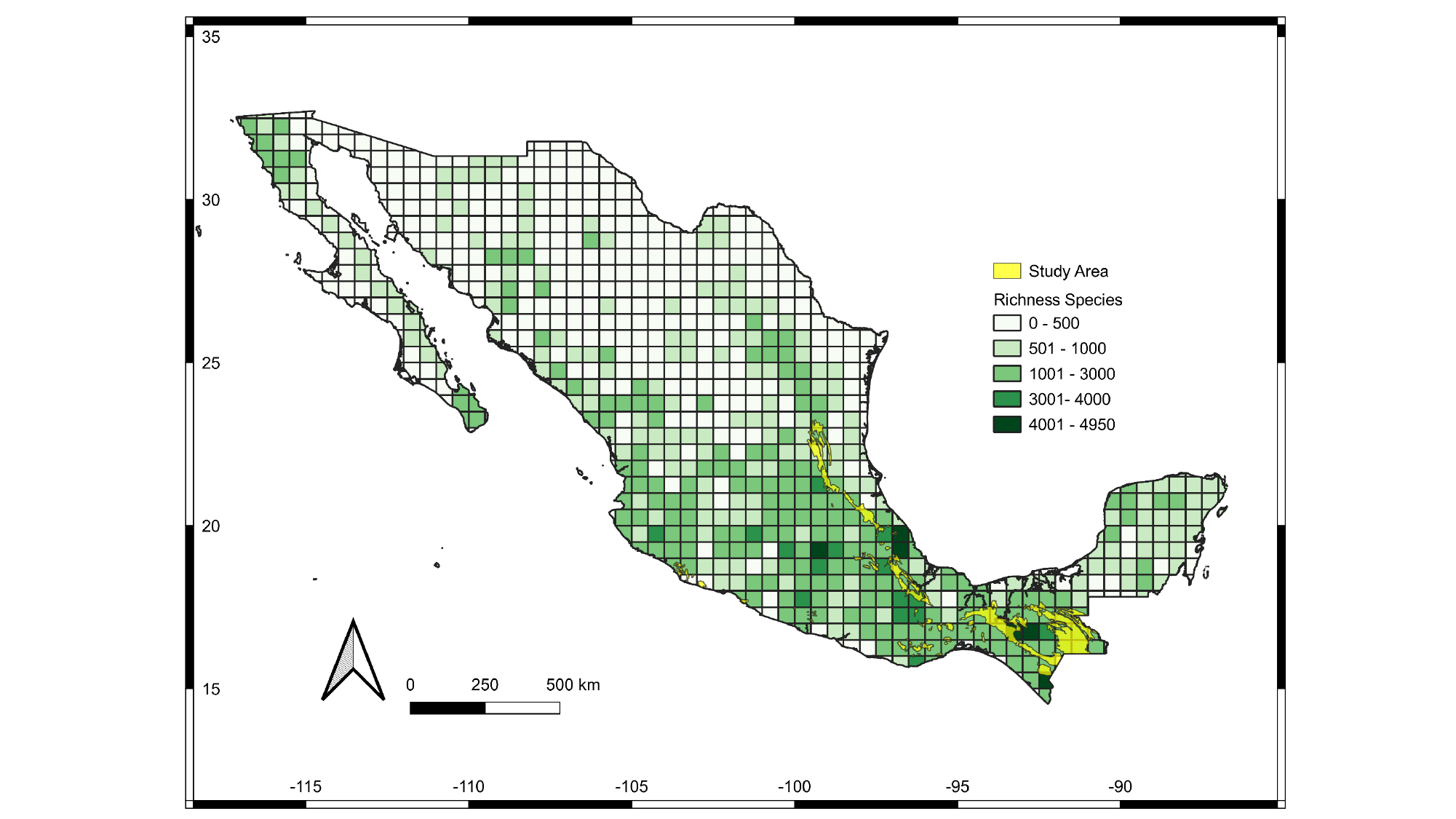
